# *araucan* regulates butterfly wing iridescence by coordinating scale structure and pigmentation

**DOI:** 10.1101/2023.11.21.568172

**Authors:** Martik Chatterjee, Cédric Finet, Kate J. Siegel, Shen Tian, Yifei Joye Zhou, Ling Sheng Loh, Xin Yi Yu, Seydeanna Delgado, Jeanne M.C. McDonald, Eirene Markenscoff-Papadimitriou, Anyi Mazo-Vargas, Antónia Monteiro, Robert D. Reed

**Affiliations:** Cornell University; Department of Biological Sciences, Faculty of Science, National University of Singapore, Singapore, 117543, Singapore; Department of Biology, Duke University, Durham, NC, 27708, USA; George Washington University; Department of Molecular Biology and Genetics, Cornell University, Ithaca, NY, 14853, USA; Department of Ecology and Evolutionary Biology, Cornell University, Ithaca, NY,14853, USA

**Keywords:** structural coloration, iridescence, butterfly wing patterns, *Iroquois*-complex, evo-devo

## Abstract

Butterfly wings exhibit a remarkable diversity of structural iridescent colors, yet the genetic regulation of iridescence remains poorly understood. Here, we show that the *Iroquois*-complex transcription factor gene *araucan* plays a role in modulating wing scale iridescence in the common buckeye butterfly, *Junonia coenia*. Using CRISPR-Cas9 knockouts, we demonstrate that loss of *araucan* function causes dorsal wing scales to shift from golden-brown to blue iridescence, and eyespot center scales to shift from saturated purple-violet to dull grey-brown. These phenotypes are associated with changes in thickness of the lower lamina in scale cells, a structural feature known to influence thin-film interference, along with reduction of pigmentation in ground scales. Together, these results identify *araucan* as a single transcription factor that determines coloration by simultaneously regulating both photonic architecture and light-absorbing pigmentation, and show that these effects can be regulated in pattern-specific manner. Our findings provide a framework for linking gene regulation to the spatially coordinated evolution of structural and pigmentary components of color.

## INTRODUCTION

Coloration in animals serves as a powerful model for understanding how gene regulatory networks give rise to complex, adaptive traits. Color is most commonly generated by light-reflecting pigments, by structural features that alter light reflectance spectra, or through a combination of both. Although structural coloration is widespread across the animal kingdom—particularly in insects—its genetic and developmental bases remain poorly understood (1). Butterfly wings, with their nanostructured scales and vivid structural organization that produce blue, purple, and green hues, offer a useful system for uncovering how genes shape the physical traits that underlie these colors.

Over the past decade advances in genomic mapping and CRISPR-Cas9 gene editing have enabled the discovery and functional validation of key genes and regulatory elements involved in butterfly wing pigmentation and patterning. Comparative studies and insights from model insects such as *Drosophila melanogaster* have further expanded the developmental genetic framework for wing pattern determination.

However, the genetic basis of structural coloration remains less well resolved, with only a handful of causative genes identified across a few butterfly species. These include *optix*, a regulator of blue iridescence in *Junonia coenia* (2,3) and silver scale development in *Speyeria mormonia* (4); *bric-a-brac* and *doublesex*, which control UV-reflecting scales in *Colias spp.* (5) and *Zerene spp.* (6), and *antennapedia*, *apterousA*, *doublesex*, *ultrabithorax*, *optix*, and *hr38*, which regulate silver reflectance in *Bicyclus anynana* (7,8). Of these examples, *optix* is the only gene thus far characterized to play a role in iridescent coloration.

Here, we investigate the role of *araucan/caupolican* (also known as *iroquois2*), an *Iroquois*-complex transcription factor gene, in the wing patterning of the common buckeye butterfly *J. coenia*. Most hexapods, including butterflies, possess two *Iroquois* genes—one orthologous to *D. melanogaster’s mirror* (*iroquois1*) and one to *araucan/caupolican* (*iroquois2*). In *D. melanogaster iroquois2* has undergone a lineage-specific duplication, giving rise to the paralogs *araucan* and *caupolican* (9–11). The three *Iroquois* genes (*araucan*, *caupolican*, and *mirror*) in *D. melanogaster* regulate multiple developmental processes, including dorsal notum patterning, longitudinal wing vein specification, and posterior wing domain (alula) identity (12–16). Recent findings suggest that *mirror* specifies a far posterior domain in butterfly wings—a role likely ancestral in broad winged insects but lost in *D. melanogaster* and other calyptrate flies due to evolutionary reductions in wing complexity (11). Despite its expression in the basal part of the wing in *Heliconius* butterflies (17), the function of *araucan/caupolican* (hereafter *araucan*) in butterflies has thus far remained untested.

In this study, we used CRISPR-Cas9 mutagenesis and gene expression analyses to investigate the role of *araucan* in wing coloration in *J. coenia*. We show that *araucan* is a regulator of both wing scale ultrastructure, including the iridescence-determining lower lamina thickness, and scale pigmentation. Our findings identify *araucan* as a novel regulator of coloration in butterflies that functions via a process of coordinated pigment and structural effects. This work provides the first functional evidence for this gene outside of *D. melanogaster*, highlighting its evolutionary flexibility and specialized role in modulating scale-based structural color traits.

## MATERIALS AND METHODS

### Identification of *Iroquois-*complex genes

We previously identified the *Iroquois*-complex genes in *J. coenia* (11). For this study, we used an updated annotation (18), which resulted in revised gene model identifiers for the *Iroquois*-complex. We identified the *J. coenia* gene *JC_g12213* as the single ortholog of *D. melanogaster araucan/caupolican* paralogs (figure S1). For clarity and consistency, we refer to *JC_g12213* as *araucan* in *J. coenia*, with the understanding that it is equally related to both the paralogs in *D. melanogaster*. The updated phylogeny and sequence information used for orthology assignments are provided in figures S1 and file S1, respectively. To ensure reproducibility and compatibility with future genome and annotation releases, we have provided both the predicted amino acid and coding DNA sequences of *araucan* in the supplementary materials (file S1).

### CRISPR-Cas9 mutagenesis of *J. coenia araucan*

To assess the function of *araucan* in *J. coenia* we generated CRISPR-Cas9 somatic mosaic deletions targeting the *araucan* coding region. We used a previous protocol where early embryos are injected with single guide RNAs (sgRNAs) and Cas9 enzyme

(19). Mutations induced in early syncytial embryos typically only affect a subset of nuclei, resulting in mosaic knockout (mKO) animals.

We designed two sgRNAs – 5’-CTGGGTACGATTTAGCAGCC-3’ and 5’-CTCCTGCTCCTTGTCCTCGT-3’ – to target the exon of *araucan* containing the conserved homeobox domain in *J. coenia* (figure S1a). Possible off target sgRNAs were ruled out by blasting against the *J. coenia* genome assembly. sgRNAs were synthesized by Integrated DNA Technologies, Inc. (Coralville, IA, USA), rehydrated in 1x Tris-EDTA buffer, and mixed with Alt-R S.p. Cas9 Nuclease/Nickase (Integrated DNA Technologies, Inc., Coralville, IA, USA) to a final concentration of 200ng/ul sgRNA and 500ng/ul Cas9.

We collected eggs by placing *Plantago lanceolata* leaves in a cage with gravid females for 1-2 hours, and then prepared eggs for microinjection by submersing them in 5% benzalkonium chloride (Sigma-Aldrich, St Louis, MO, USA) for 30s followed by a wash with distilled water. Prepared eggs were then lined up on double sided tape on glass slides and sgRNA and Cas9 mixture was injected into the eggs using 0.5-mm borosilicate needle (Sutter Instruments, Novato, CA, USA) and a PLI-100 Picoinjector (Harvard Apparatus, Holliston, MA, USA). Injections were completed within 2-3hrs after oviposition, and larvae from injected eggs were raised on artificial diet at 27°C, 70% humidity, and a 16hr : 8hr light : dark cycle. Adults were frozen upon emergence, and phenotyped and imaged using a Keyence VHX-7000 microscope.

### Genotyping CRISPR mutants

We genotyped *J. coenia* mosaic knockout (mKO) mutants to confirm Cas9-induced deletions in the *araucan* gene. Mutants “mKO A” and “mKO 3” were selected based on their blue cover scale phenotypes (figure S2), while “mKO M” was chosen for its symmetrically disrupted eyespot phenotype (figure S4). We extracted DNA (OMEGA BIO-TEK, Norcross, GA, USA) from the thoracic wing muscles of butterflies that showed obvious wing pattern anomalies compared to wild types. We used PCR to amplify DNA from around the potential cut sites using the primers 5’-ATACTAAACCCTGCACCCGC-3’ and 5’-ACGTACTCACAAGCGAACGT-3’. We ran amplified DNA from wild type and mutant individuals on an agarose gel and purified (Monarch DNA Extraction and Purification kit, new England Biolabs, MA, USA) and cloned the PCR products into JM109 competent cells using pGEM®-T Easy Vector Systems (Promega Corporation, Madison, WI, USA). Transformed cells were amplified and Sanger sequenced at the Genomics Facility, Cornell Institute of Biotechnology, Ithaca, NY, USA.

### Micro-spectrophotometry of wing scales

We mounted individual wing scales (n = 3) from wild-type and *araucan* mutant *J. coenia* butterflies on glass slides and measured their normal-incidence reflectance spectra using a mercury-xenon light source (Thorlabs, New Jersey, USA) connected to a uSight-2000-Ni micro-spectrophotometer (Technospex Pte. Ltd., Singapore). We used a Nikon TU Plan Fluor 100× objective (numerical aperture = 0.9) to collect spectra and calibrated the setup with a dielectric mirror. For each measurement, we averaged 10 scans with an integration time of 100 ms. We recorded three measurements from different locations on each of three scales per genotype and averaged them to obtain a representative reflectance spectrum.

To measure absorbance, we used the same optical setup with a 20× objective (NA = 0.5). We mounted individual scales (n = 5) submerged in clove oil (Hayashi Pure Chemical Ind., Ltd.) between a slide and coverslip to match the refractive index of chitin and used a clove oil-covered transparent area of the slide as a reference. For each genotype, we measured three locations on each of five scales and averaged the data to generate absorbance spectra.

We performed all spectral analysis and visualization using the pavo 2 package in R

(20). To test for significant differences in spectral values between wild-type and *araucan* mutant scales, we conducted independent two-sample t-tests at each 50 nm increment from 350 to 700 nm. We applied the Benjamini-Hochberg False Discovery Rate (FDR) method to adjust p-values for multiple comparisons and considered adjusted p-values < 0.05 as significant.

### Scanning electron microscope imaging of wing scales

We used scanning electron microscopy (SEM) to assess structural differences between wild-type and *araucan* mutant scales. We mounted individual scales (n = 5) from mutant mosaic clones and contrasting wild-type regions on carbon tape and sputter-coated them with platinum (JEOL JFC-1600) for 30 s at 30 mA. We imaged the samples using a FEI Versa 3D with the following parameters: voltage 10 kV, current 32 pA. To obtain transverse-sections of wing scales, we used Focused Ion Beam (FIB) milling with the gallium ion beam of the FEI Versa 3D, applying a beam voltage of 8 kV, beam current of 25 pA, and a tilt of 52°. We performed the FIB-SEM in the Electron Microscopy Facility, National University of Singapore.

We obtained the transverse section of the scales shown in Figure 1c by cryo-fracturing the scales following a previously described protocol (21). We then mounted whole and cryo-fractured scales of interest on carbon tape and sputter-coated them with an 8 nm layer of gold using a Cressington 208HR Sputter Coater. We acquired the image on an FEI Teneo LV SEM using the Everhard Thornley detector and beam parameters of 5.00 kV/6.3 pA with a 10 µs dwell time. We used cryo-fracturing only to generate the image used in figure 1c. All measurements for laminar thickness reported were done on scales that were milled using FIB and imaged using SEM.

**Figure 1.**
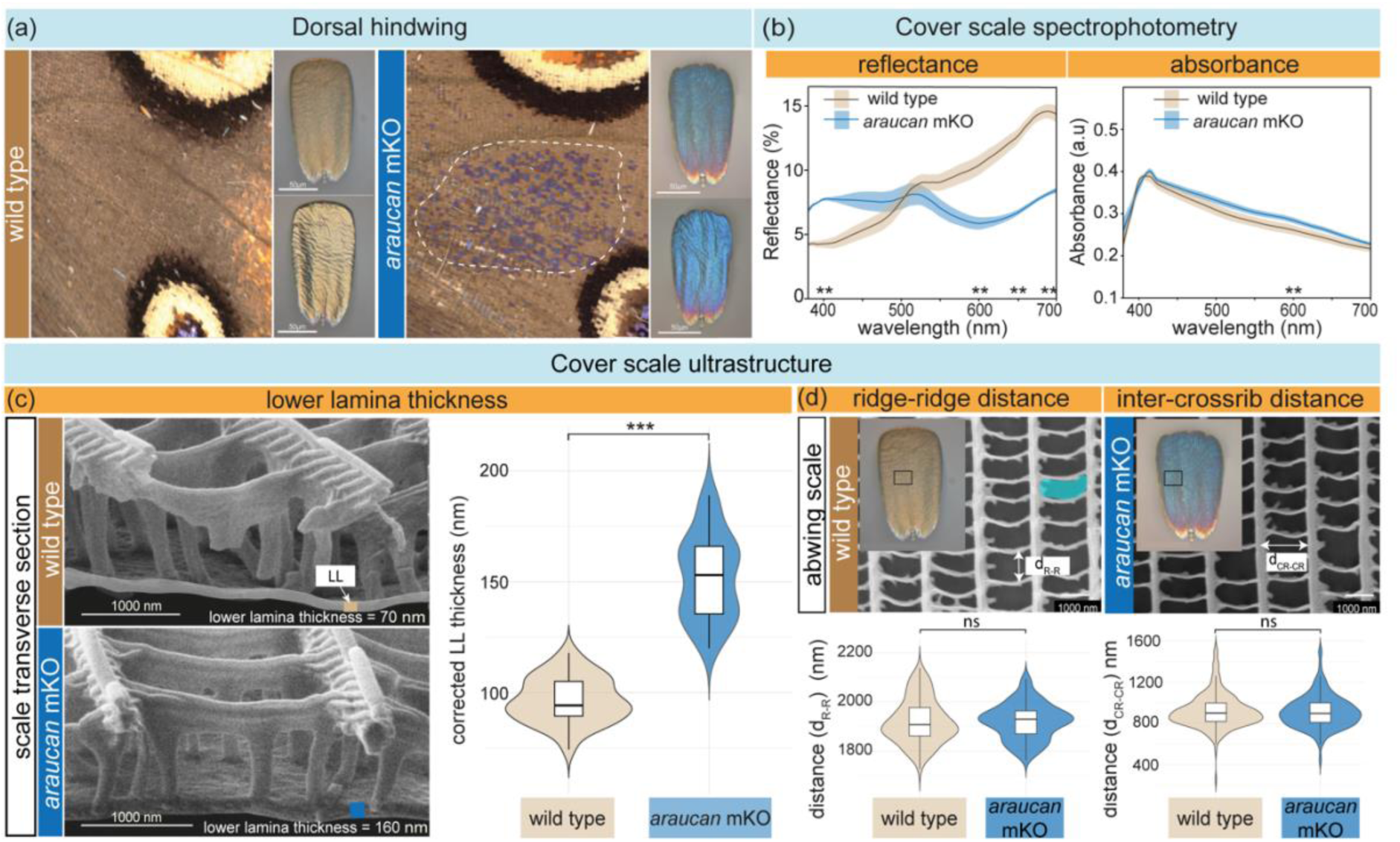
*araucan* knockouts cause blue iridescence in dorsal hindwings by increasing lower lamina thickness of cover scales. (a) *araucan* mKO produces patches of blue iridescent scales in the dorsal hindwing. *Left*: Wild-type wing showing uniform golden-brown color. Adjacent are magnified views of individual cover scales from the same wing, with the abaxial surface (*top*) and adaxial surface (*bottom*) shown. Right: *araucan* mKO hindwing showing a mosaic patch of blue iridescent scales (*dashed line*). Adjacent are magnified views of mutant cover scales, revealing blue iridesence on both the abaxial (*top*) and adaxial (*bottom*) surfaces. Scale bar = 50 μm. Additional images of contralateral wings showing mutant blue iridescent scales are in figures S2 and S3. (b) Reflectance and absorbance spectra of cover scales confirm that the observed color shift is structural, not pigment-based. Solid lines indicate mean spectra; shaded regions represent standard deviation; asterisks mark wavelengths with significant differences between groups (n_reflectance_ = 3; n_absorbance_ = 5; \*\**p-adj* < 0.01; table S2). (c) *Left:* SEM images of transverse sections from isolated cover scales, prepared by cryo-fracturing, show that wild-type scales (*top*) have a thinner lower lamina (LL; *grey shaded region*) compared to mutant scales (*bottom*). *Right*: Violin-box plot quantifies lower lamina thickness from FIB-milled isolated cover scales (n = 5; \*\*\**p* < 2 × 10⁻¹⁶; tables 1, S1). (d) Additional cover scale measurements: ridge-to-ridge spacing (d_R-R_, *top left*), distance between crossribs (d_CR-CR_, *top right*), and area of the windows (*teal*) - through which incident light goes into the scale (*bottom*, ns – not significant; tables 1, S1).

### Measurement of scale ultrastructure and statistical analysis

We quantified scale ultrastructure by measuring three parameters: lower lamina thickness (LL), inter-ridge distance (d_RR_), and crossrib (d_CR-CR_) spacing (figures 1c-d). We performed all measurements in Fiji using the line tool (22). Lower lamina thickness was corrected for tilted perspective (measured thickness/sin 52°) (23). For each phenotype (i.e. wild type or *araucan* mutant), we analyzed five individual scales per wing region and scale type. From each scale, we recorded 10 measurements of lower lamina thickness, 25 measurements of inter-ridge distance, and 50 measurements of crossrib spacing. We collected measurements from cover and ground scales in the forewing eyespot center, and from cover scales in the dorsal non-eyespot region.

We analyzed scale ultrastructure data using linear mixed-effects (LME) models implemented in the R package nlme version 3.1 (24) which accounts for hierarchical data structures. For each geometric parameter (lower lamina thickness, inter-ridge distance, and crossrib spacing), we treated scale type as a fixed effect and included scale identity nested within individual butterfly as a random effect. To address heteroscedasticity among scale types, we applied the varIdent() function. We selected the best-fitting model based on the Akaike Information Criterion (AIC). We conducted pairwise comparisons of group means using the multcomp R package. We report p-values and 95% confidence intervals for all pairwise comparisons in Tables 1 and 2. A summary of the fixed effects from the linear mixed-effects models, including degrees of freedom, F-values, and p-values, is provided in Supplementary Tables S2 and S3. Raw measurement data are also included to support independent reanalysis (see Data Availability).

**Table 1.** Comparison of wild-type and *araucan* mutant non-eyespot cover scale ultrastructural features.

| Scale type | Factor | Comparison | Estimate | Std error | Z-value | Pr(>\|z\|) | 95% CI |
| --- | --- | --- | --- | --- | --- | --- | --- |
| cover | Lower-lamina thickness | WT - mutant == 0 | -55.41 | 2.93 | -18.89 | <2e-16<br>*** | [-61.16 ; -49.66] |
| cover | Distance ridge-ridge | WT - mutant == 0 | 0.54 | 0.54 | 10.13 | 0.05 | [-19.32 ; 20.39] |
| cover | Distance crossrib-crossrib | WT - mutant == 0 | 8.61 | 14.25 | 0.6 | 0.54 | [-19.31 ; 36.54] |

**Table 2.** Comparison of wild-type and *araucan* mutant cover and ground scale ultrastructural features collected from eyespot center.

| Scale type | Factor | Comparison | Estimate | Std error | Z-value | Pr(>\|z\|) | 95% CI |
| --- | --- | --- | --- | --- | --- | --- | --- |
| cover | Lower-lamina thickness | WT - mutant == 0 | 38.92 | 2.61 | 14.89 | <2e-16<br>*** | [33.79 ; 44.04] |
| cover | Distance ridge-ridge | WT - mutant == 0 | 317.75 | 6.83 | 46.51 | <2e-16<br>*** | [304.36 ; 331.14] |
| cover | Distance crossrib-crossrib | WT - mutant == 0 | -256.73 | 24.15 | -10.63 | <2e-16<br>*** | [-304.06 ; -209.40] |
| ground | Lower-lamina thickness | WT - mutant == 0 | 38.46 | 2.45 | 15.72 | <2e-16<br>*** | [33.67 ; 43.25] |
| ground | Distance ridge-ridge | WT - mutant == 0 | 282.4 | 6.87 | 41.08 | <2e-16<br>*** | [268.93 ; 295.88] |
| ground | Distance crossrib-crossrib | WT - mutant == 0 | -32.52 | 7.49 | -4.34 | 1.42e-05<br>*** | [-47.20 ; -17.83] |

### Hybridization chain reaction fluorescent *in situ* hybridization of *araucan*

We designed a hybridization chain reaction (HCR) probe set targeting the coding sequence of *J. coenia araucan* using 19 pairs of 25 bp probes (50 bp per site) spanning the annotated transcript (file S2). We ordered the probes as oligonucleotides from Genewiz (Azenta Life Sciences, Morrisville, NC). We confirmed probe specificity using BLAST against the most recent *J. coenia* genome assembly, ensuring no off-target binding sites.

For HCR experiments, we dissected last instar larval imaginal discs, as well as pupal wings at 13% (first day post-pupation), 25% (second day post-pupation) and 45% (third day post-pupation) of pupal development. We used *araucan* probes paired with B1 initiators and visualized them with Alexa Fluor 647-labeled hairpins. As a positive control, we used *spalt* probes labelled with B3 initiators (file S2) and visualized them in the 546 nm channel. For figure S6 we performed HCR of *optix* with B5 initiator (file S2). We performed co-detection of *araucan* and spalt expression in wings from one side of each individual, while using the contralateral wings as negative controls. In these controls, we omitted the probes but followed all other HCR steps to assess background fluorescence. We performed the HCR following the protocol described by Bruce et al (dx.doi.org/10.17504/protocols.io.bunznvf6). To capture for pattern specific expressions across the entire wing tissue we used a Keyence BZ-X800 fluorescence microscope in the laboratory of Dr. Anyi Mazo-Vargas (Department of Biological Sciences, Duke University).

## RESULTS

### *araucan* knockouts result in iridescent blue scales via laminar thickening

We injected 758 eggs with Cas9 and *araucan* sgRNAs, of which 179 hatched and 117 survived to adulthood. Twenty-two of the adults that emerged had wing color mutations. We observed two distinct classes of mutations, both of which involved iridescent coloration: (1) blue iridescence in mutant clones across the dorsal wing surface (n = 13), and (2) a shift from purple-violet to grey-brown specifically in eyespot centers (n = 9). Genotyping confirmed that individuals from both mutant classes had lesions at the sgRNA sites in the *araucan* coding region (figure S1c).

The thirteen individuals that showed mosaic clones of blue iridescence showed the phenotype exclusively on dorsal wing surfaces (figure 1a; hindwings: figure S2; forewings: figure. S3). Since iridescent blue colored scales can be artificially selected for in populations of *J. coenia* (3), we assessed individuals from our wild type colony for similar iridescent scales to rule out the possibility of phenotypes being due to natural variation. Out of the individuals randomly observed (n = 32), we did not observe any blue iridescent scales on the dorsal surface of the hind wings, which supports that the blue iridescence in the mKOs is a result of knockdown of *araucan* (chi-squared test, *p* < 0.005). This conclusion was also supported by the fact that individuals with iridescent patches of scales presented this phenotype asymmetrically (figures S2 and S3), consistent with cell lineage mosaicism (2), whereas natural variation in iridescence is typically symmetrical.

The shift from golden-brown scales in the wild type to vivid blue in mutants was quantified using reflectance spectrophotometry. Mutant scales exhibited increased reflectance in the ultraviolet and blue wavelengths (< 500 nm), while wild-type scales reflected more light in longer wavelengths (> 500 nm), consistent with their golden-brown appearance (figure 1b). To determine whether this observed color change was structural or pigmentary in origin, we immersed individual scales in oil, which matches the refractive index of chitin and thereby suppresses structural coloration. As in previous studies, absorbance spectra collected under these refractive index-matched conditions provide a measure of the light transmitted through the scale and served as a proxy for pigment content, as differences in pigment type and concentration result in distinct patterns of wavelength-specific light absorption (3,7,21,25–28). We observed minimal differences in absorbance of visible wavelengths between mutant and wild-type scales (figure 1b; table S2), indicating that the shift in coloration is primarily due to structural changes rather than differences in pigmentation.

To investigate the ultrastructural basis of the iridescent blue coloration in *araucan* mutant scales, we measured the thickness of the lower lamina—a key structural determinant of scale color in butterflies of the *Junonia* genus, where a thicker lower lamina causes a shift towards bluer hues (3). We quantified laminar thickness in both wild-type brown scales and blue mutant scales to assess whether this relationship held true in our *araucan* knockouts. SEM imaging after FIB milling to get the transverse section of scales revealed that mutant cover scales had significantly thicker lower laminae compared to wild type (figure 1c; LME model, AIC-selected best fit, *p* < 2 × 10⁻¹⁶; tables 1, S1). The mean corrected lower lamina thickness in wild-type scales was 96.7 ± 10.2 nm, while mutant scales averaged 152.0 ± 18.9 nm, supporting the hypothesis that the observed color shift is structurally mediated.

To assess whether additional ultrastructural features contribute to the iridescent color shift observed in *araucan* mutants we quantified ridge spacing and crossrib distances in cover scales. These geometric features determine the size of the scale “windows” that determine the amount of incident light that interacts with underlying scale nanostructures (figure 1d). Mean ridge-to-ridge spacing was comparable between wild-type brown and mutant blue scales (WT: 1921 ± 90 nm; mutant: 1921 ± 73 nm; figure 1d; tables 1, S1), and crossrib distance was also unchanged (WT: 923 ± 167 nm; mutant: 914 ± 160 nm; figure 1d; tables 1, S1). These results suggest that the shift to blue structural coloration in the *araucan* mutants is not due to changes in ridge or crossrib geometry, but instead is primarily driven by increased lower lamina thickness.

### *araucan* knockouts eliminate purple-violet iridescence in forewing eyespot centers

Nine of the recovered *araucan* mKOs exhibited a striking phenotype affecting the scales in the forewing eyespot centers. In wild-type *J. coenia*, these eyespot centers exhibit a vivid purple-violet iridescence (figures 2a and S4). In contrast, *araucan* mutants lacked this strong purple-violet coloration, which was replaced by a muted grey-brown hue, occasionally with faint traces of pink or violet iridescence (figures 2a and S4). The loss of purple-violet iridescence was asymmetrical in all but two individuals (figure S4), validating the mosaic nature of the knockout phenotype. Notably, eyespot centers on the hindwings were unaffected.

**Figure 2.**
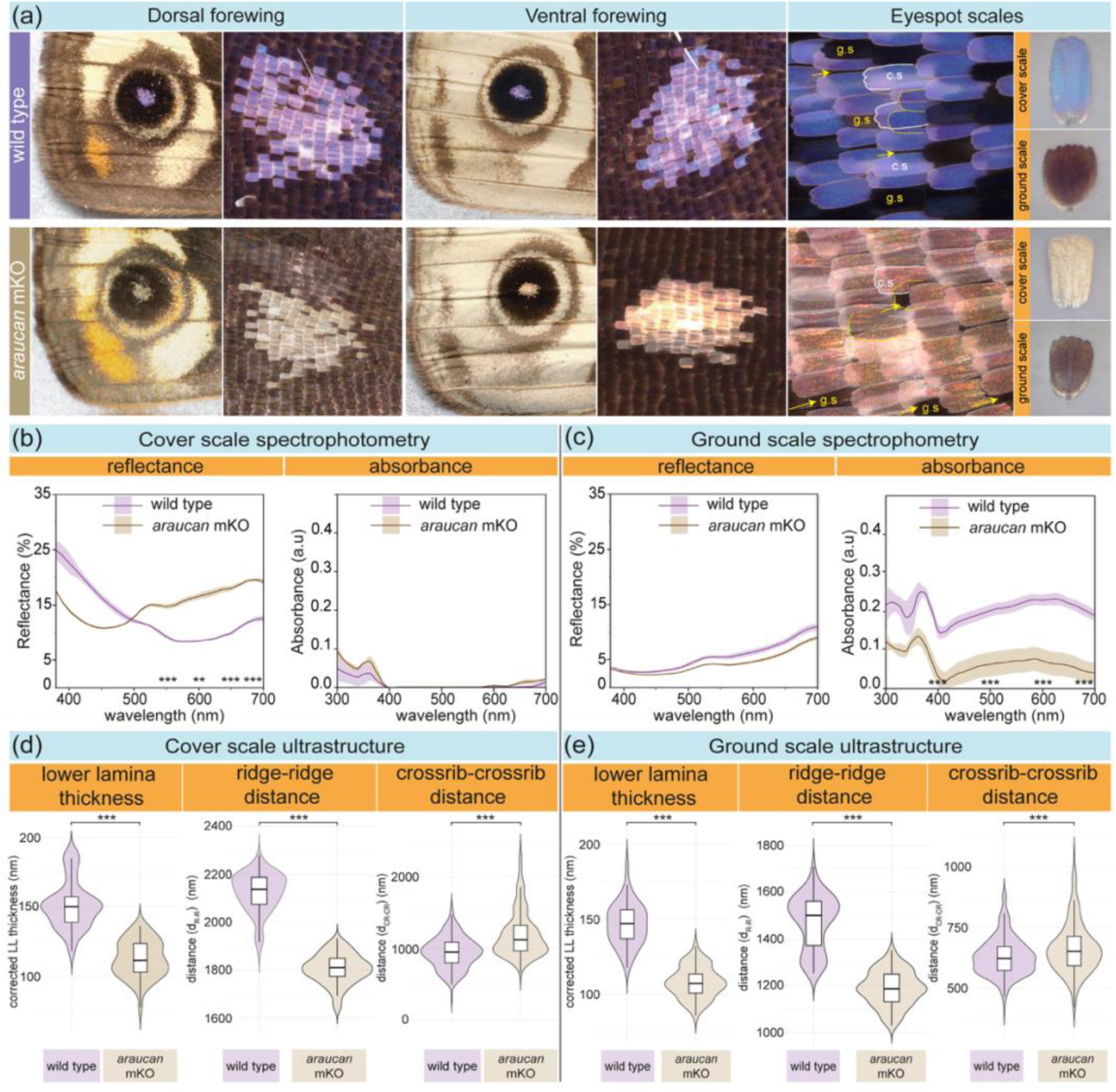
*araucan* knockouts diminish purple-violet iridesence by altering cover scale reflectance, ground scale pigmentation, and the ultrastructure of both scale types in forewing eyespot centers. (a) On both dorsal (*left*) and ventral (*middle*) surfaces, forewing eyespot centers of wild-type individuals (*top*) exhibit distinct purple-violet iridescence, whereas *araucan* knockouts (*bottom*) show muted grey-brown scales (figure S4). Magnified images of the eyespot center (*right*) highlight the stacking arrangement of cover (c.s; *white border*) and ground scales (g.s; *yellow arrows where exposed and yellow dashed boundaries where overlaid by cover scales*). Overlaid ground scales highlight saturation of overall color (*yellow dashed boundary*). Abwing surfaces of isolated cover and ground scales (*far right*) of wild-type (*top*) and mutant (*bottom*) scales. (b) Spectrophotometry of isolated cover scales and (c) ground scales from eyespot centers in wild-type and mutant forewings. Reflectance spectra (*left*) and absorbance spectra (*right*). Solid lines represent mean spectra; shaded areas indicate standard deviation; asterisks denote wavelengths with significant differences between groups (n_reflectance_ = 3; n_absorbance_ = 5; ***p-adj < 0.001, **p-adj < 0.01; Table S4). (d–e) Violin-box plots quantifying ultrastructural features in cover scales (d) and ground scales (e) (n = 5; ***p < 2 × 10⁻¹⁶; Tables 2, S3).

The iridescent purple-violet coloration of *J. coenia* eyespot centers arises from the combined optical properties of both cover scales and underlying ground scales. The ground scales enhance the overall saturation of the reflected color when stacked beneath the more translucent cover scales (highlighted by yellow dashed outlines in figure 2a). Because this mechanism is consistent with observations in other butterfly species exhibiting purple-violet coloration (29,30), we characterized the optical properties of each scale type within the eyespot center. Specifically, we measured reflectance spectra and refractive index-matched absorbance spectra of both cover and ground scales in the forewing eyespot center to identify which scale type and optical mechanism was contributing to the observed color shift.

Spectral analysis showed that both scale types, cover and ground, were altered in the *araucan* mutants differently, resulting in the observed color shift. Mutant cover scales exhibited increased reflectance above 500 nm, corresponding to a duller grey-brown coloration, whereas wild-type cover scales reflected more light in the ultraviolet to violet wavelengths (380–450 nm), producing the observed purple-violet hue (figures 2a, b). In contrast, reflectance measurements of ground scales revealed no significant differences between wild-type and mutant individuals (figure 2c). To assess pigment contributions independent of structural effects, we measured absorbance in a refractive index–matched medium. Under these conditions, both wild-type and mutant cover scales showed minimal light absorption in the ultraviolet and red regions, with no significant differences between groups, indicating that most incident light was transmitted through the cover scales (figure 2b). However, mutant ground scales showed a different pattern across the ultraviolet and visible spectrum (300–700 nm) where they exhibited significantly lower absorbance compared to wild-type scales, suggesting reduced light-absorbing pigmentation in the mutants (figure 2c).

To identify structural components in the scales that might have been altered, we characterized lamina thickness, ridge spacing, and crossrib distances in both cover and ground scales. We found that the lower lamina was significantly thicker in the eyespot scales of wild-type individuals compared to *araucan* mutants in both scale types. In cover scales, the lower lamina thickness averaged 151 ± 16.9 nm in wild types and 112 ± 13.1 nm in the mutants (LME model, AIC-selected best fit, *p* < 2e-16; figure 2d; tables 2 and S3). A similar pattern was observed in ground scales (146 ± 13.9 nm in wild types vs 108 ± 10.3 nm in mutants, *p* < 2e-16; figure 2e; tables 2 and S3).

Ridge spacing was significantly narrower in mutants versus wild-type scales, for both scale types. In cover scales, ridge spacing was 2123 ± 82.9 nm in wild types and 1805 ± 62.9 nm in mutants (*p <* 2e-16; figure 2d; tables 2 and S3), and in ground scales, 1470 ± 116 nm in wild types versus 1188 ± 74 in mutants (*p* < 2e-16 figure 2e; tables 2 and S3). In contrast, the distance between crossribs was significantly greater in *araucan* mutants for both scale types. In cover scales, mutants had crossrib spacing of 1202 ± 322 nm compared to 946 ± 217 in wild types (*p* < 2e-16; figure 2d; tables 2 and S3); and in ground scales, mutants had 663 ± 103 nm of spacing vs 631 ± 79.3 nm in wild types (*p =* 1.42e-05; figure 2e; tables 2 and S3). Together, these optical and structural analyses reveal that *araucan* knockouts disrupt the pigmentation of ground scales and the nanostructural organization of both cover and ground scales in the eyespot centers. In combination, these changes likely underlie the loss of saturated purple-violet iridescence and its replacement with muted hues in *araucan* mutants.

### Dynamic spatial and temporal expression of *araucan* in the developing wing

Our mKOs demonstrate that *araucan* is necessary for normal structural coloration of *J. coenia* wing scales and appears to play a role in determining scale structure itself.

Accordingly, we hypothesized that we should observe *araucan* expression in wings during scale morphogenesis. Because *Iroquois*-complex genes—including *araucan*—exhibit pattern-specific expression in the *D. melanogaster* wing disc (12,13), we further sought to ask whether *araucan* is expressed in a pattern-specific manner in the developing *J. coenia* wings, including in regions affected in mutants (e.g., eyespot centers), or in other defined wing domains across development. To characterize spatial transcriptomic patterns, we performed HCR *in situ* hybridization for *araucan* mRNA at key developmental stages: (1) in last-instar wing discs, immediately during and after the transcription factor Spalt defines future eyespot domains (31–33); and (2) in pupal wings at two time points −13% pupal development (PD), when eyespot centers have been established, and 25% PD, when scale precursors differentiate into scale cells (34).

In last-instar discs, *araucan* transcripts are enriched in the posterior wing domain (vannus) bounded by the 2A vein (figure 3a). Additional signal appears in an anterior–proximal region and in enlarged cells arrayed along the trachea that mark future veins and the wing margin (figures 3a and S5b). At ∼13% PD, the posterior-domain expression persists and *araucan* is detected in large cells interspersed along the veins of both forewings and hindwings (figure 3b). By ∼25% PD, posterior-domain signal contracts to a narrow stripe flanking the 2A vein, while expression becomes broadly peppered across both wings without aligning to obvious adult color elements (figure 3c). This expression, along with the expression in the veins, persists into later pupal development (figure S5e). Across stages co-stained with the eyespot marker *spalt*, we did not detect *araucan* expression obviously upregulated in eyespots (figure S5), despite peppered expression spanning those regions in the stages sampled after 13 % pupal development (figure 3c).

**Figure 3.**
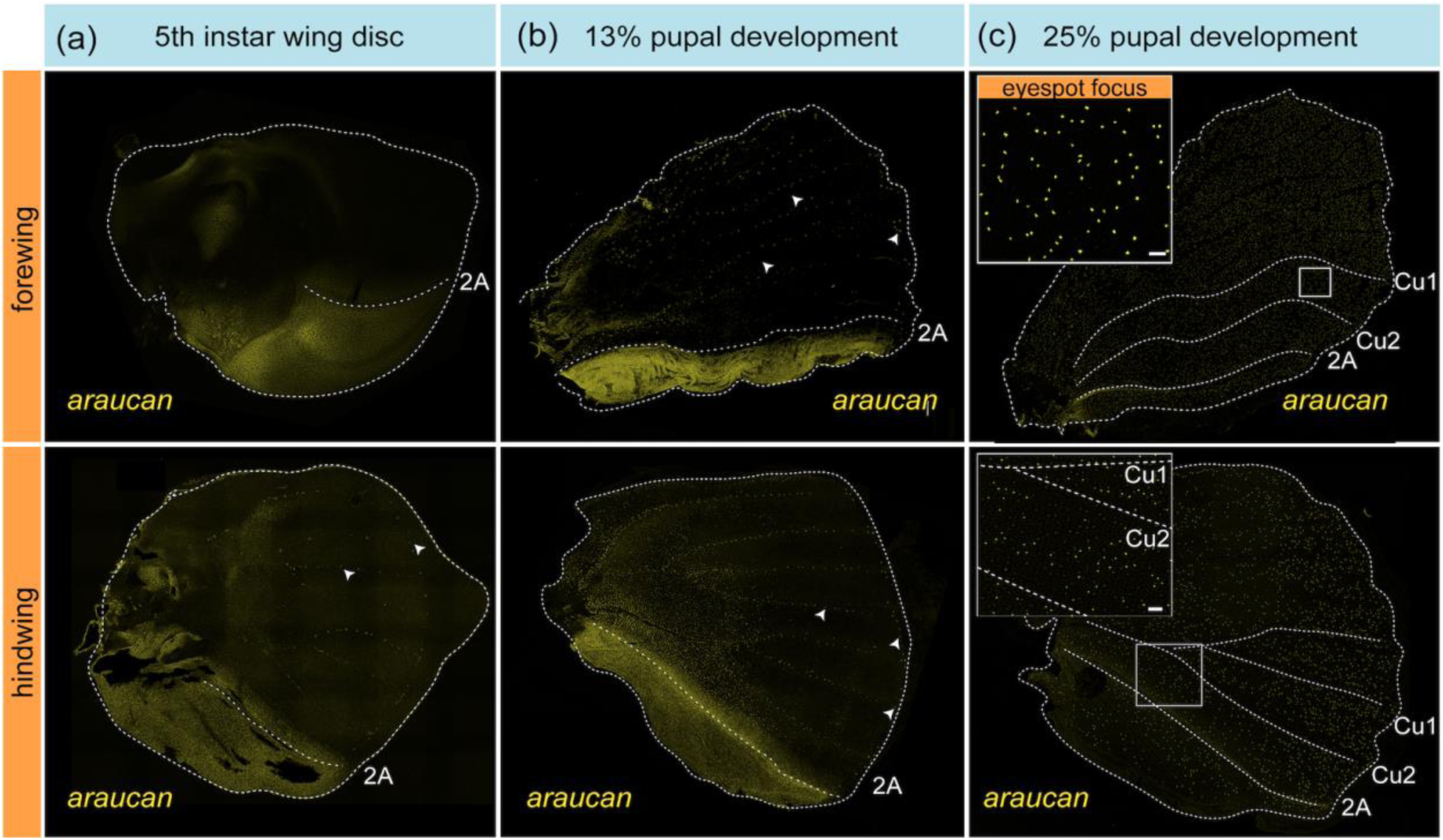
Spatiotemporal dynamics of *araucan* transcripts in late larval and early pupal wings. (a–b) In the 5th-instar larva (a) and ∼13% pupal development (b), *araucan* signal (yellow) is enriched in the far-posterior vannus domain, bounded anteriorly by the 2A vein in both wings. Signal is also detected in cells along pre-vein tracheae and at the wing margin (white arrows). (c) By ∼25% pupal development, *araucan* persists along the 2A vein and appears in scattered cells across the wing blade in a pattern-nonspecific manner (yellow dots). In the forewing (*top*), this includes the eyespot focus (*inset*). In the hindwing, this recaptures the mosaic clones seen in the mKOs (*inset*). [Scale bar = 50 μm]

Together, these data show that *araucan* transitions from early domain- and vein-associated patterns to a later dispersed distribution in developing scale cells. This later expression pattern mirrors the spatially scattered scale iridescence phenotypes in mKO patches and is consistent with a role for *araucan* in scale-cell differentiation and morphogenesis occurring at the assayed stage.

## DISCUSSION

### Region specific control of iridescence by *araucan*

In this study, we investigated the role of the *Iroquois*-complex gene *araucan* in butterfly wing patterning. Our findings reveal that *araucan* is a key regulator of structural coloration in *J. coenia*, modulating iridescence in two spatially and functionally distinct ways: (1) by suppressing blue iridescence across the dorsal wing surface, and (2) by promoting purple-violet iridescence in forewing eyespot center.

Structural coloration in butterfly wings arises from complex interactions between incident light and the ultrastructure of wing scales, as reviewed by Thayer and Patel (35). One well-characterized mechanism is thin-film interference, in which the lower lamina of the scale functions as a reflective surface, producing a range of iridescent hues depending on its thickness and the angle of incident light (29,35–37). Previous work in *Junonia spp*. has shown that the thickness of the lower lamina within individual cover scales reliably predicts spectral reflectance: thin laminae generate golden-brown hues as seen in wild type *J. coenia*, whereas thicker laminae produce blue or teal hues as seen in *optix* knockouts of *J. coenia* or species in the genus that naturally exhibit blue-green iridescence (3). In our *araucan* knockouts, we observed an increase in lower lamina thickness in dorsal cover scales, resulting in a shift from brown to blue structural coloration. This optical shift occurred without detectable changes in scale pigmentation, or other scales indicating that *araucan* suppresses iridescence in non-eyespot regions by constraining laminar thickness.

In contrast, the eyespot centers of wild-type *J. coenia* display purple-violet iridescence that emerges from the combined optical effects of stacked cover and ground scales as shown in other nymphalid butterflies (29,30,36). The ultrastructure and pigmentation of the cover scales modulate the amount of incident light reflected or transmitted, while those of the underlying ground scales influence the extent of light that is scattered back. Together, these layers generate interference patterns that either enhance or dampen the resulting color signal. In *araucan* mutants, we observed increased laminar thickness in both cover and ground scales, accompanied by a marked reduction in ground scale pigmentation. We infer that the loss of pigment decreases light absorbance, thereby increasing backscatter, which—when combined with structural modifications in the cover scales—produces a desaturated, muted tan-grey hue in place of the saturated purple-violet seen in wild-type eyespots. This enhanced backscatter likely contributes to the shimmering or hazy appearance of iridescence observed in some mutant individuals (figure S4). In addition, differences in ridge spacing and crossrib distances may further influence optical properties, either directly or by influencing pigmentation levels as seen before (27). Together, these findings demonstrate that *araucan* regulates multiple dimensions of scale morphology and pigmentation that collectively shape the optical characteristics of forewing eyespot center coloration.

The opposing effects of *araucan* in eyespot versus non-eyespot scales underscore a broader principle in butterfly wing development: regulatory genes often function in a region-specific, context-dependent manner. The divergent roles of *araucan* reinforce the idea that eyespots represent modular, semi-independent developmental units capable of evolving specialized scale properties—including iridescence—without altering patterning in adjacent regions. This spatial modularity enables flexible evolutionary innovation within the wing, raising important questions about the regulatory architecture controlling *araucan* expression across the wing.

### Gene regulatory dynamics of *araucan*

We were intrigued to observe that *araucan* had a very dynamic pattern of transcription over developmental time, ranging from vein- and posterior domain-related patterns in early development, to a whole-wing scattered pattern later in pupal development that is consistent with iridescence phenotypes. The early posterior domain expression pattern was of particular interest because it resembles the expression of *mirror*–the sister gene of *araucan* which has been shown to play a role in posterior domain determination (11). Curiously, we did not observe any domain-related or otherwise *mirror*-like phenotypes in our *araucan* mutants and it therefore remains unclear whether this posterior *araucan* expression pattern has any functional relevance. It is interesting, however, to speculate whether the parallel expression patterns of *araucan* and *mirror* may reflect ancestral regulatory architecture that predates the divergence of these genes. In any case, the complex and dynamic expression patterns of *araucan* over developmental time suggest this gene is likely responding to a varied progression of upstream trans-regulatory signals.

Our CRISPR mKOs and expression data raise the question of where *araucan* sits within the gene regulatory networks that pattern the wing and specify scale-cell fates. One of the two major mKO classes we observe was a blue shift in the iridescence of dorsal wing scales, consistent with increased lower-lamina thickness in cover scales. A very similar effect was previously reported for *optix* mKOs in *J. coenia*. Indeed, the blue shift effect observed in *araucan* knockouts could reasonably be described as a subset of *optix* knockout effects. Although we did not detect marked co-expression of *optix* and *araucan* in the same cells at the timepoints we sampled (figure S6), this does not rule out participation in a shared pathway: transient or low-abundance transcripts, development staging, or non-cell-autonomous interactions could all evade overlap in our datasets. We therefore propose two working models that explain the similar effects of these genes: (1) a shared pathway in which *optix* and *araucan* act in series or converge on common ultrastructural effectors to modulate lower-lamina thickness (yielding similar blue-shift phenotypes), and (2) parallel pathways that independently alter scale ultrastructure and pigment deposition but produce convergent phenotypes. Future work should assess to what degree *optix* and *araucan* may work together to determine scale cell structural coloration.

In addition to structural blue shifting of dorsal wing scales, *araucan* had a different, yet highly specific effect on the structural and pigmentary coloration of eyespot centers.

How might *araucan* be deployed for such a different, spatially restricted eyespot function? In *J. coenia*, mKO of *spalt* eliminates all eyespots (38), consistent with *spalt* acting as a master regulator of eyespot formation. Because the *araucan* phenotype tracks *spalt* expression patterns, one plausible model is that *spalt* functions upstream of *araucan* in eyespot centers. The absence of detectable *spalt–araucan* co-expression in our HCR in situs does not rule this out: *araucan* eyespot transcription may be below our detection threshold or expressed at later stages of wing development, which makes sense for a gene that regulates terminal stages of scale maturation. A similar detection–phenotype mismatch is familiar from other wing transcription factors (e.g., *optix*), where *in situ* hybridization patterns from earlier stages of development (figure S6) contrast with broader phenotypic effects of knockouts. This all suggests that gene expression patterns during later stages of development, which are challenging to image for technical reasons, are important for structural and pigmentary color determination (2). The forewing-specific *araucan* eyespot phenotype also lead us to speculate that *Ultrabithorax*, a selector gene for hindwing identity (39–42), may play an upstream role in regulating *araucan* in this context.

## Conclusion

Overall, our results identify *araucan* as a region-specific regulator of structural coloration in *J. coenia*, suppressing blue iridescence across the dorsal wing while promoting saturated purple–violet at the forewing eyespot focus. These effects map to coordinated changes in scale ultrastructure—especially lower-lamina thickness—and, in eyespots, to coupled shifts in optical layering with ground-scale pigmentation. Together, the data reveal how a single transcription factor can differentially tune photonic architecture across the wing, and reinforce eyespots as modular, context-dependent developmental units. While the precise placement of *araucan* within color pattern gene regulatory networks remains unresolved, our findings establish a framework for dissecting the regulatory logic that links gene activity to the coordinated evolution of cell structure and pigment synthesis.

## Data availability

The raw SEM images used for ultrastructural measurements, associated data tables, raw spectral data from scale micro-spectrophotometry, genome assembly and annotation files, are available through Dryad (DOI: DOI: 10.5061/dryad.rfj6q57n6).

## SUPPLEMENTARY INFORMATION

**Figure S1.**
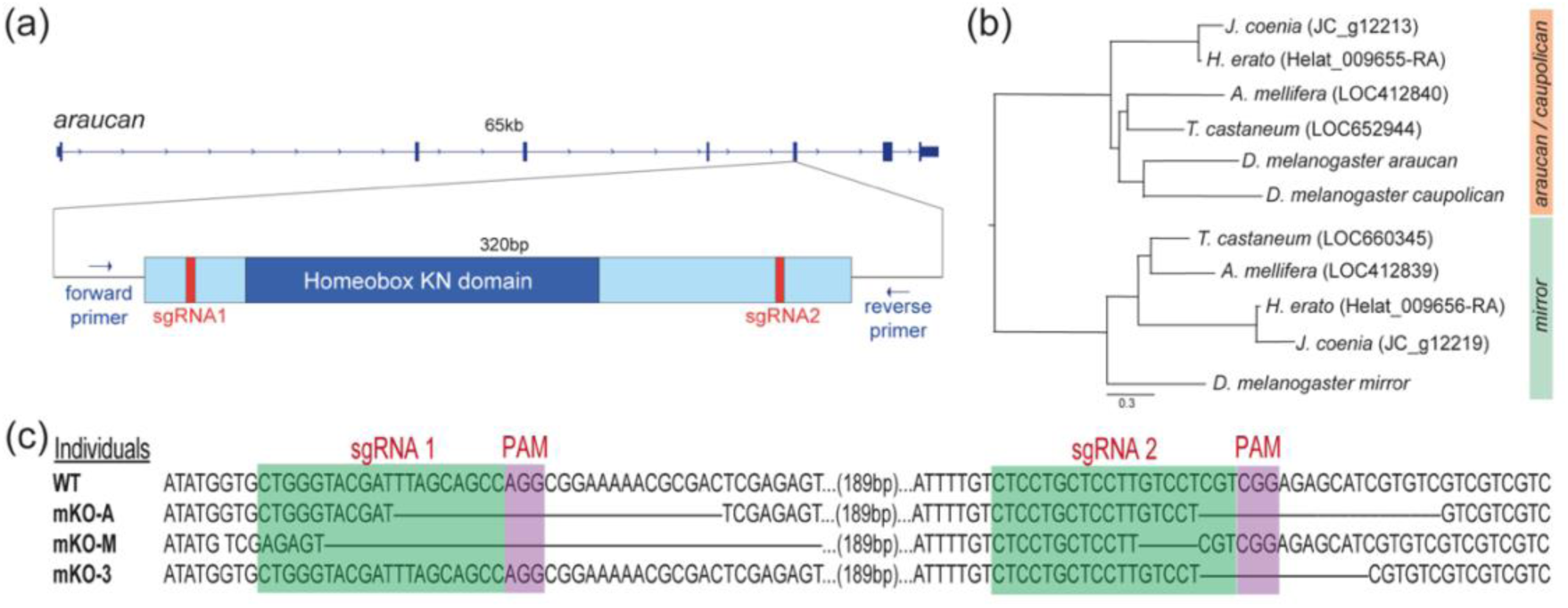
(a) Map of the *araucan* locus in *J. coenia,* annotated with the location of sgRNAs. The primers shown were used to validate deletions in mutants. Refer to Materials and Methods for the sequences of sgRNAs and genotyping primers. (b) Maximum likelihood phylogeny of *J. coenia*, *Heliconius erato lativitta*, *Tribolium castaneum*, *Apis mellifera*, and *D. melanogaster Iroquois* complex genes show that *JC_g12213* is the *araucan*/*caupolican* ortholog in *J. coenia* and *JC_g12219* is the ortholog of *mirror*. Re-analysed already published data (Chatterjee et al. 2025), using new gene model identifiers. (c) CRISPR-Cas9 targeting of *araucan* produces deletions at the sgRNA sites. Wild type and mutant sequences are shown. Green boxes indicate sgRNA binding sites. Purple boxes denote PAM sites Knockouts mKO “A” and mKO “3” show blue dorsal iridescence phenotypes (Figure S2). mKO M has eyespot focus phenotype (Figure S2).

**Figure S2.**
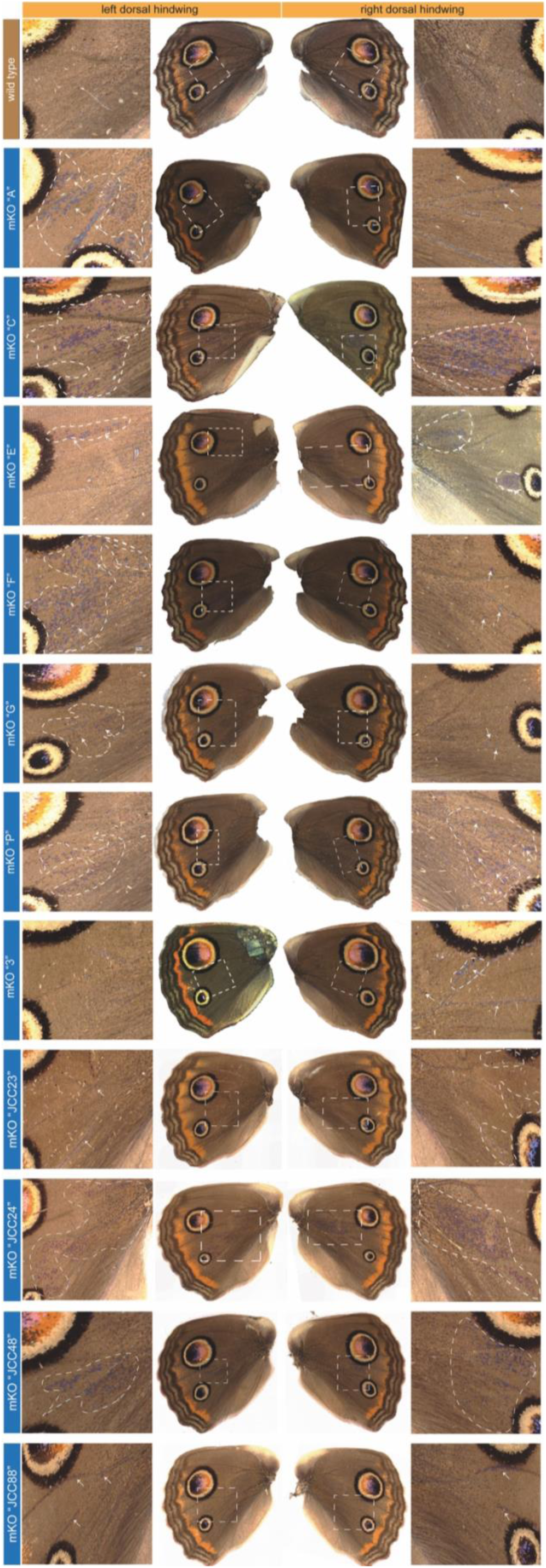
*araucan* mKOs displaying mutant patches (dashed lines) of iridescent blue scales (arrows) in hindwings. Contrasting lateral wings from individual butterflies are shown to highlight the asymmetrical nature of the mosaic phenotypes.

**Figure S3.**
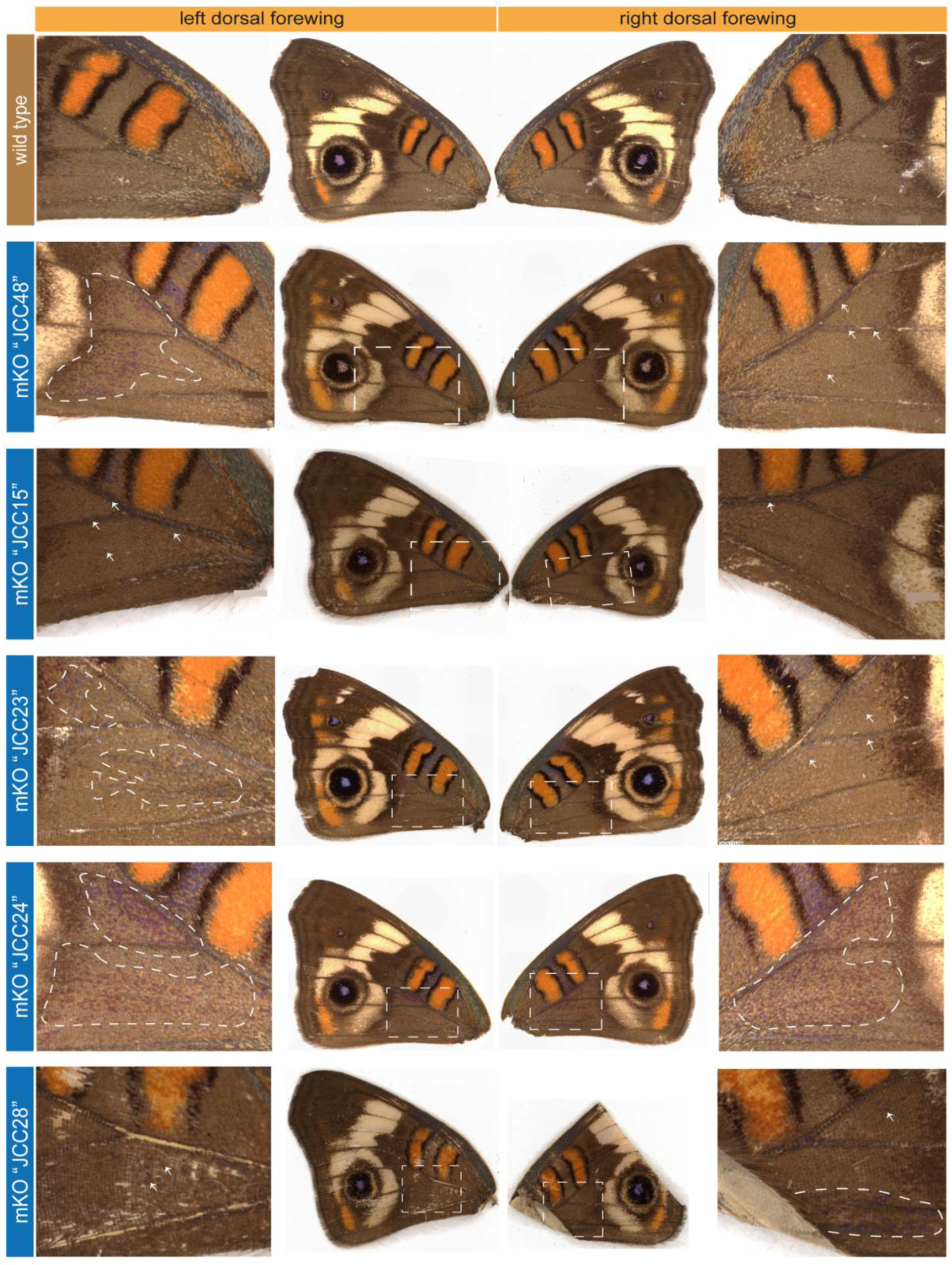
*araucan* mKOs displaying mutant patches (dashed lines) of iridescent blue scales (arrows) in a subset of mutant forewings. Contrasting lateral wings from individual butterflies are shown to highlight the asymmetrical nature of the mosaic phenotypes.

**Figure S4.**
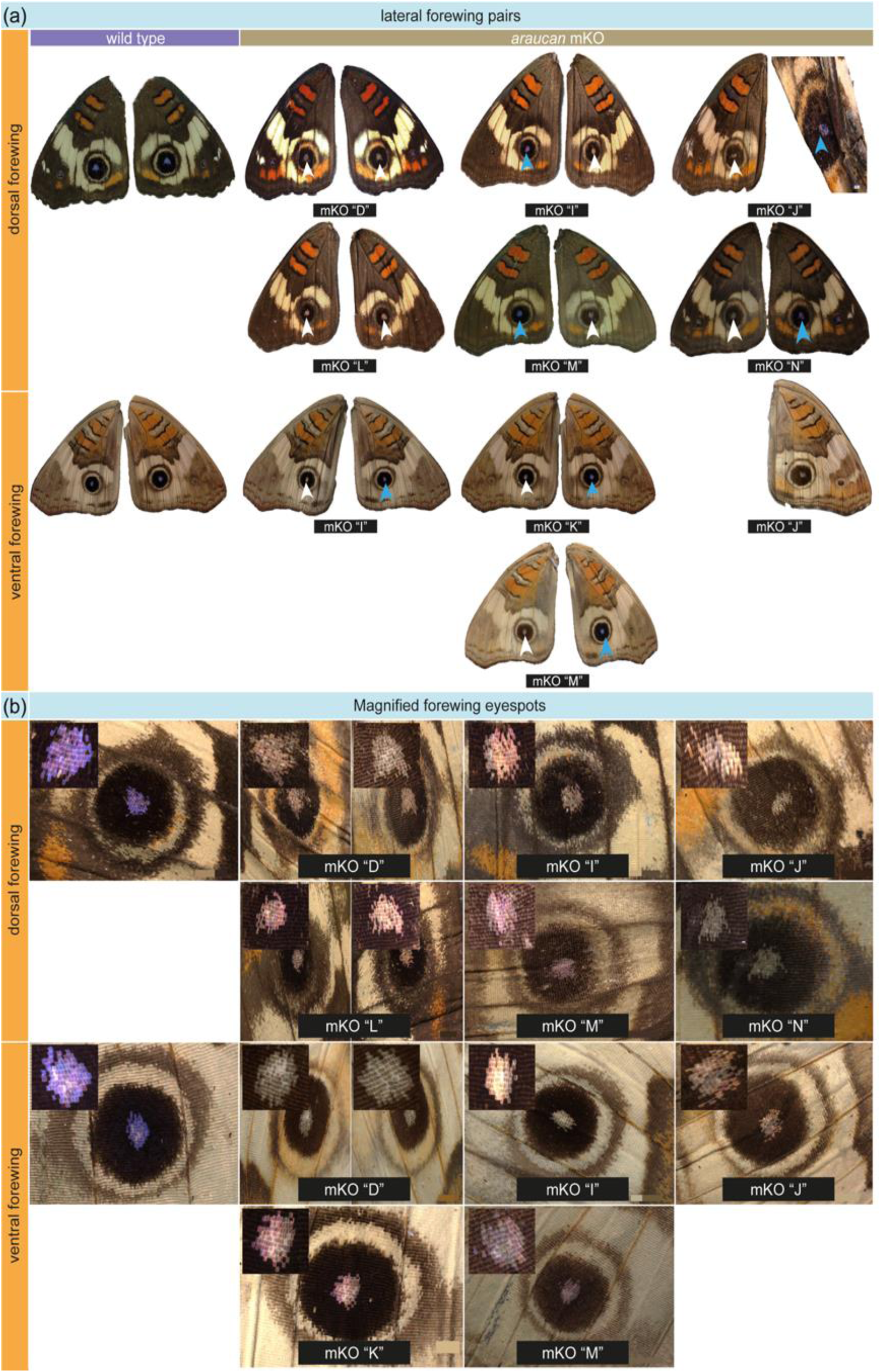
Loss of purple-violet iridescence in forewing eyespot foci following *araucan* mosaic knockout (mKO). (a) Contralateral forewings from representative individuals showing the elimination of purple-violet iridescence in dorsal (*top*) and ventral (*bottom*) eyespot foci. In most cases, both forewings exhibited the mutant phenotype, with exceptions noted in specimens “mKO D” and “mKO L.” Blue arrows indicate wild-type foci with saturated iridescent coloration; white arrows indicate mutant foci displaying altered or absent iridescence. (b) Enlarged views of eyespot foci highlighting the phenotypic effects of *araucan* mKOs. Mutant scales show a range of altered hues, from muted grey to shimmering pink-purple, compared to the vivid purple-violet observed in wild type. Insets show scale-level differences in the dorsal (*top*) and ventral (*bottom*) surfaces.

**Figure S5.**
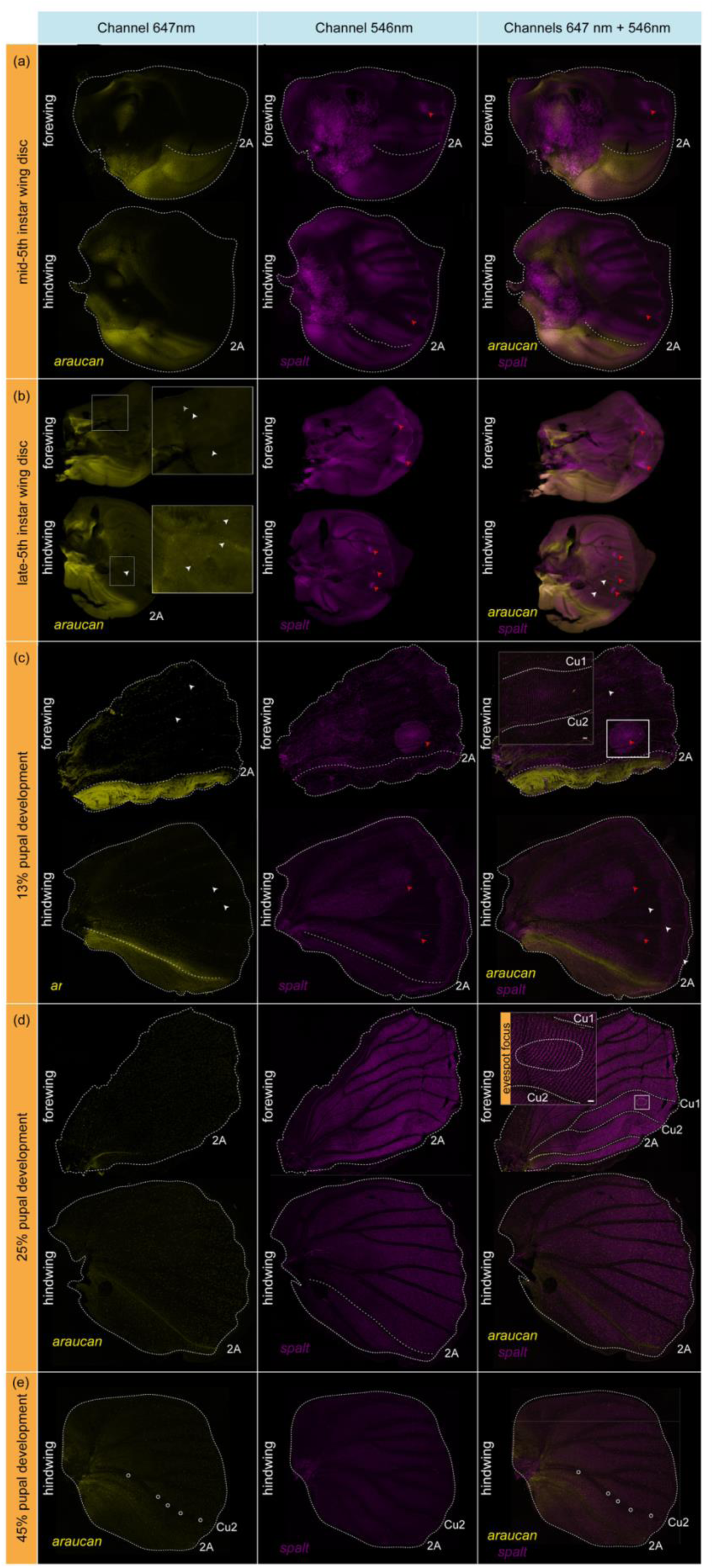
Comparative expression of *araucan* and *spalt* during wing development. Each panel shows three columns: left, *araucan* mRNA (yellow); middle, *spalt* mRNA (magenta); right, merge. (a–b) 5th-instar larval wings: *araucan* is enriched in the posterior domain and in discrete cells along pre-vein tracheae (white arrows). *spalt* marks future eyespot domains (red arrows). No overlap between *araucan* and *spalt* is detected at this stage. (c) ∼13% pupal wings: the larval pattern persists—*araucan* expression remains to the posterior wing and large cells along the veins (white arrows), while *spalt* marks eyespot fields (red arrows) with no detectable co-expression in the eyespot focus (inset). (d) ∼25% pupal wings: *spalt* is broadly expressed across scale-forming cells, whereas *araucan* is retained along the posterior/2A domain and scattered cells across the wing; with no distinct difference in the focus (inset). (e) ∼45% pupal development: Only hindwing showing expression patterns of *araucan* and *spalt* similar to earlier stage with no overlap. White circles highlight large, interspersed cells along the Cu2 vein. [scale bar = 50 μm]

**Figure S6.**
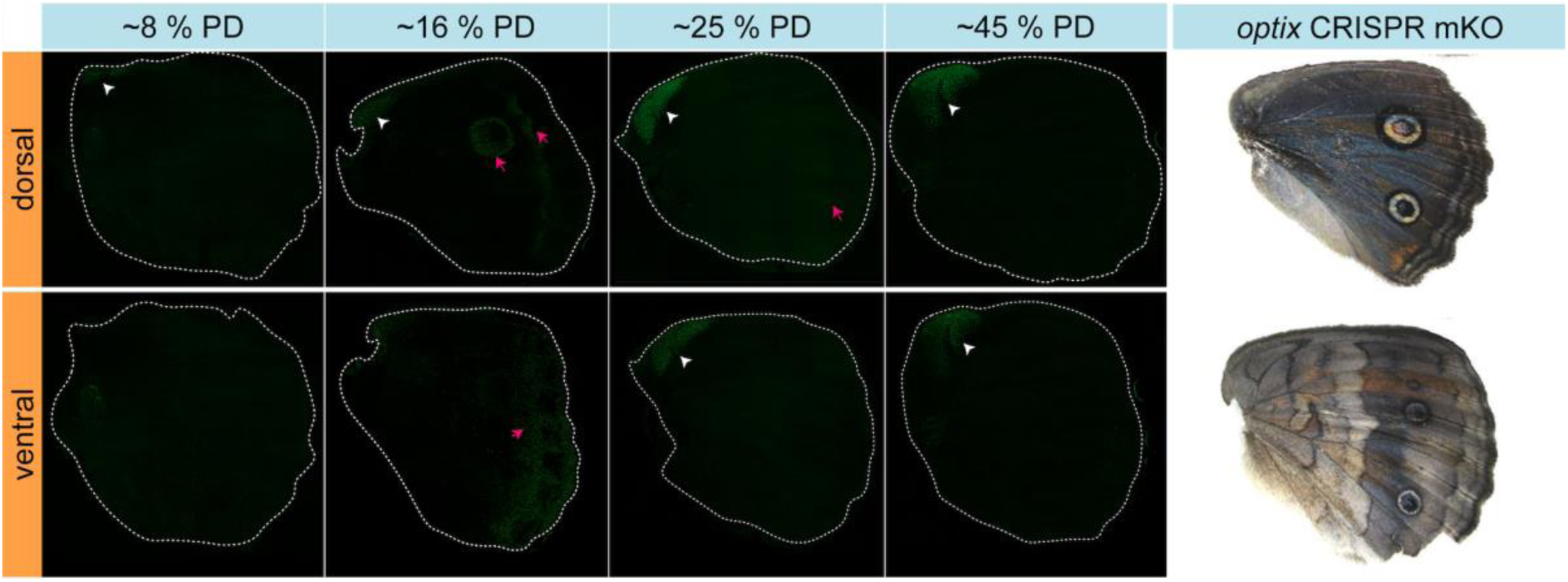
Optix mRNA expression in early to mid-pupal development (PD) captures only a subset of large CRISPR knockout effects in adults. Strong *optix* expression in the wing conjugation scales across development (white arrowhead). Pattern specific expression in the eyespots and parafocal elements (magenta arrow) peaks at 16% pupal development and gradually lowers in 25% and 43% development. *Optix* mKO phenotypes in *J. coenia* adults results in large effects across the wing (2), despite early pupal spatial expression. *Optix* mKO CRISPR images taken from Zhang and Mazo-Vargas et. al (2).

**Table S1.**
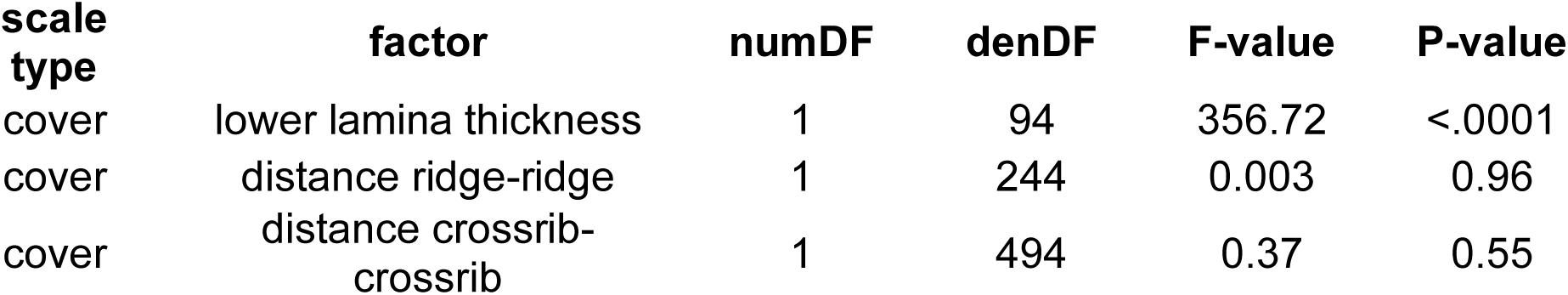
Comparison of wild-type and *araucan* mutant non-eyespot cover scale ultrastructural features.

**Table S2.**
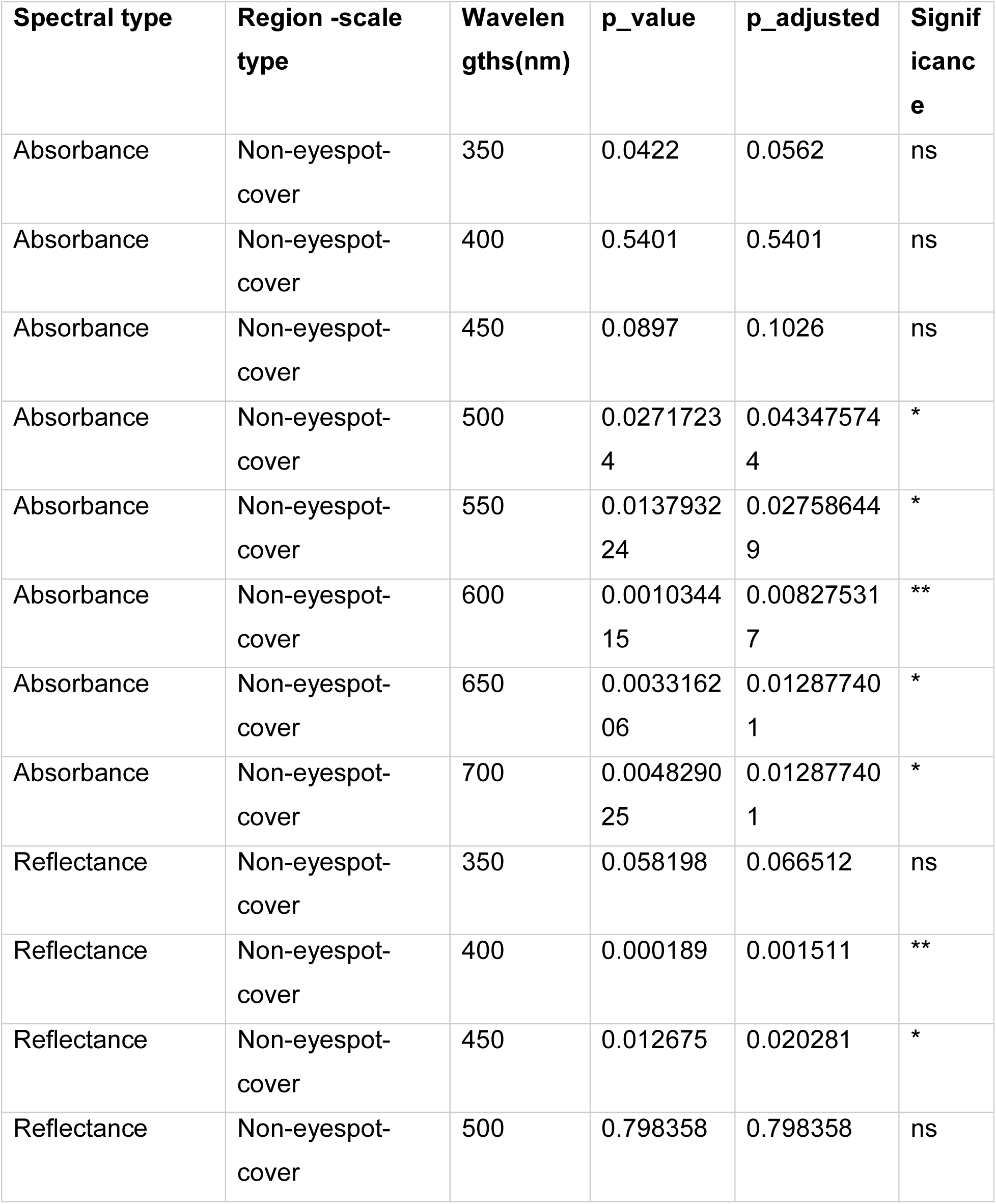
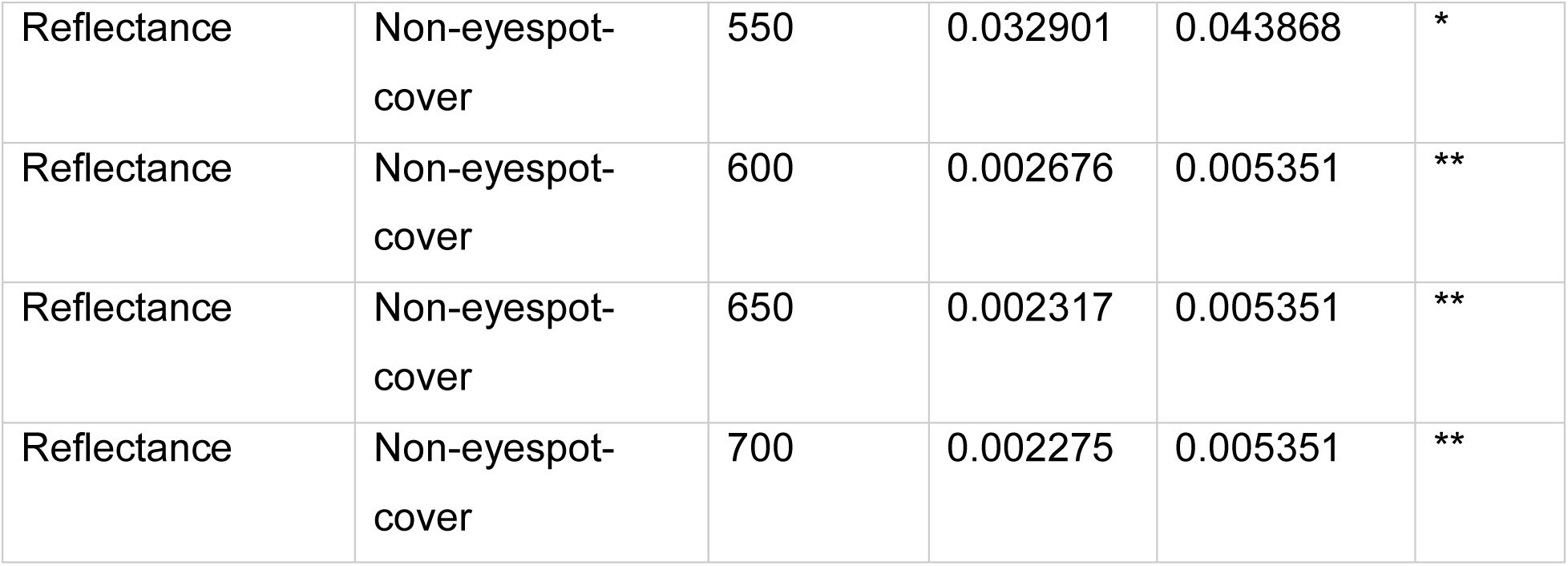
Significance table of reflectance and absorbance spectra measured at different wavelengths.

**Table S3.** Comparison of wild-type and *araucan* mutant cover and ground scale ultrastructural features collected from eyespot center.

| scale type | factor | numDF | denDF | F-value | P-value |
| --- | --- | --- | --- | --- | --- |
| cover | lower lamina thickness | 1 | 94 | 221.66 | <.0001 |
| cover | distance ridge-ridge | 1 | 244 | 2163.07 | <.0001 |
| cover | distance crossrib-crossrib | 1 | 494 | 113.02 | <.0001 |
| ground | lower lamina thickness | 1 | 98 | 247.29 | <.0001 |
| ground | distance ridge-ridge | 1 | 244 | 1687.48 | <.0001 |
| ground | distance crossrib-crossrib | 1 | 494 | 18.84 | <.0001 |

**Table S4.** Significance table of reflectance and absorbance spectra measured at different wavelengths.

| Spectral type | Region - scale type | Wavelengths( nm) | p_value | p_adjust ed | Significan ce |
| --- | --- | --- | --- | --- | --- |
| Reflectance | Eyespot-ground | 350 | 0.425298076 | 0.4252981 | ns |
| Reflectance | Eyespot-ground | 400 | 0.153736635 | 0.175699 | ns |
| Reflectance | Eyespot-ground | 450 | 0.080597206 | 0.1289555 | ns |
| Reflectance | Eyespot-ground | 500 | 0.035926762 | 0.0958047 | ns |
| Reflectance | Eyespot-ground | 550 | 0.098515912 | 0.1313545 | ns |
| Reflectance | Eyespot-ground | 600 | 0.024533269 | 0.0958047 | ns |
| Reflectance | Eyespot-ground | 650 | 0.002884728 | 0.0230778 | * |
| Reflectance | Eyespot-ground | 700 | 0.05407291 | 0.1081458 | ns |
| Reflectance | Eyespot-cover | 350 | 0.227171385 | 0.2596244 | ns |
| Reflectance | Eyespot-cover | 400 | 0.015502189 | 0.0206696 | * |
| Reflectance | Eyespot-cover | 450 | 0.00821814 | 0.013149 | * |
| Reflectance | Eyespot-cover | 500 | 0.676972861 | 0.6769729 | ns |
| Reflectance | Eyespot-cover | 550 | 0.000250499 | 0.0006858 | *** |
| Reflectance | Eyespot-cover | 600 | 0.0028632<br>43 | 0.005726<br>5 | ** |
| Reflectance | Eyespot-cover | 650 | 0.0002571<br>88 | 0.000685<br>8 | *** |
| Reflectance | Eyespot-cover | 700 | 8.9404E-05 | 0.000685<br>8 | *** |
| Absorbance | Eyespot-ground | 350 | 0.3086519<br>17 | 0.308651<br>9 | ns |
| Absorbance | Eyespot-ground | 400 | 8.29698E-05 | 0.000167<br>3 | *** |
| Absorbance | Eyespot-ground | 450 | 0.0001068<br>84 | 0.000171 | *** |
| Absorbance | Eyespot-ground | 500 | 0.0001584<br>22 | 0.000181<br>1 | *** |
| Absorbance | Eyespot-ground | 550 | 8.3648E-05 | 0.000167<br>3 | *** |
| Absorbance | Eyespot-ground | 600 | 5.60723E-05 | 0.000167<br>3 | *** |
| Absorbance | Eyespot-ground | 650 | 0.0001479<br>18 | 0.000181<br>1 | *** |
| Absorbance | Eyespot-ground | 700 | 1.30136E-05 | 0.000104<br>1 | *** |

## FILES

**File S1.** mRNA and amino acid sequences of genes used in this study, along with their *D. melanogaster* orthologs. These sequences were used to construct the gene family phylogeny shown in figure S1b and to design probes for HCR *insitu* hybridization.

***File S2.*** *Sequences of araucan, spalt* and *optix HCR-probes used for in situ hybridization*.

